# CryoEM Structure of CtBP2 Confirms Tetrameric Architecture

**DOI:** 10.1101/2020.04.06.027573

**Authors:** Anne M. Jecrois, M. Michael Dcona, Xiaovan Deng, Dipankar Bandyopadhyay, Steven R. Grossman, Celia A. Schiffer, William E. Royer

**Affiliations:** Department of Biochemistry and Molecular Pharmacology, University of Massachusetts Medical School, Worcester, MA 01605; Department of Internal Medicine, Virginia Commonwealth University, Richmond, VA 23298; Department of Biostatistics, Virginia Commonwealth University, Richmond, VA 23298; Massey Cancer Center, Virginia Commonwealth University, Richmond, VA 23298

**Keywords:** C-terminal Binding-Proteins (CtBPs), Transcription regulation, Cancer, TIAM1. Metastasis

## Abstract

C-terminal binding proteins 1 and 2 (CtBP1 and CtBP2) are transcriptional regulators that activate or repress many genes involved in cellular development, apoptosis and metastasis. CtBP proteins are activated under hypoxic conditions where NAD(H) levels tend to be higher. NADH-dependent activation of CtBP2 has direct implication in multiple types of cancers and poor patient prognosis. Previous studies have proposed dimeric CtBP as the relevant oligomeric state, however our studies with multi-angle light scattering have shown that the primary effect of NADH binding is to promote the assembly of two CtBP dimers into tetramers. Here, we present the cryoEM structures of two different constructs of CtBP2 corroborating that the native state of CtBP2 in the presence of NADH is indeed tetrameric. The physiological relevance of tetrameric CtBP2 was tested in HCT116; CtBP2 −/− cells transfected with tetramer destabilizing_mutants. Mutants that inhibit tetramer formation show a decrease in expression of the CtBP transcriptional target TIAM1 and exhibit a decrease in the ability to promote cell migration. Together with our cryoEM studies, these results highlight the tetramer as the functional oligomeric form of CtBP2.

## Introduction

C-terminal Binding proteins 1 and 2 (CtBP1 and CtBP2) are co-transcriptional factors that regulate important genes in cell fate. CtBP1 was first identified as an interacting partner of the Adenovirus 2/5 E1A protein (Boyd et al., 1993); binding occurred at the C-terminal region of E1A, resulting in the name of CtBP. CtBPs can act as both activators and repressors of transcription through their interactions with multiple transcription factors and chromatin modifier enzymes (Kuppuswamy et al., 2008). Although CtBP1 and CtBP2 share over 80% amino acid sequence identity, their functions are both unique and redundantly overlapping within the cell (Hildebrand & Soriano, 2002) (Chinnadurai, 2007). Unlike CtBP1, CtBP2 has a nuclear localization signal (NLS) at its N-terminus domain suggesting a more critical role for the latter in transcription (Hildebrand & Soriano, 2002; Ma et al., 2020). The transcriptional activity of CtBPs can confer resistance to apoptosis and promote metastasis and oncogenesis depending on their interacting partners (Chinnadurai, 2009). CtBPs may be activated under conditions of hypoxia where the NADH level is elevated in the cell, which has direct implication in various forms of cancer (Di et al., 2013). CtBP expression has been observed to be higher in colorectal cancer, melanoma, metastatic prostate cancer, ovarian cancer and breast cancer (Barroilhet et al., 2013; Deng et al., 2013; Wang et al., 2012). CtBPs promote tumorigenesis by enhancing epithelial-mesenchymal transition (EMT) and conferring resistance to apoptosis by regulating the expression of genes such as *TIAM1* and *Bik,* respectively (Grooteclaes et al., 2003) (Ma et al., 2020; Paliwal et al., 2012). Furthermore, elevated levels of CtBP in tumor tissue are correlated with poorer survival in breast cancer, ovarian cancer and hepatocellular carcinoma (Zheng et al., 2015; Chawla et al., 2019). The substantial data correlating CtBP with cancer progression implicates CtBP as a potential drug target.

Oligomerization is essential for CtBP transcriptional activity, with CtBP forming dimers (Kumar et al., 2002; Nardini et al., 2003) and higher order structures (Bellesis et al., 2018; Madison et al., 2013). Binding of NAD(H) promotes oligomerization of CtBP2, which is required for transcriptional activities (Kumar et al., 2002; Zhang et al., 2002). In conditions where the level of NAD(H) is low, CtBP2 is mostly dimeric (Bellesis et al., 2018). Increasing the level of NADH in the cell promotes oligomerization of CtBP2. The activated oligomeric form of CtBP can then associate with other transcriptional coactivators and enzymes to form the CtBP-mediated repression complex (Shi et al., 2003). Although previous studies have proposed dimeric CtBP as the relevant oligomeric state (Nardini et al., 2003; Thio et al., 2004; Nardini et al., 2009; Bi et al., 2018; Mani-Telang et al., 2007; Dcona et al., 2019), our studies with multi-angle light scattering and sitedirected mutagenesis have shown that the primary effect of NADH binding is to promote the assembly of two CtBP dimers into tetramers (Bellesis et al., 2018). This was further supported by the observation that CtBP1 and CtBP2 exhibit similar tetrameric assemblies within distinct crystal lattices used for structure determination (Hilbert et al., 2014), resulting in a tetramer model for CtBP.

The goal of the present study is to determine the structure of CtBP2 in solution. Solving the structure without the influence of a crystalline lattice will permit us to investigate the importance of the tetrameric assemblage for transcriptional activity in cellular function. We have determined the solution structures of CtBP2 by cryoEM at an average resolution of 3.6Å, for the minimal dehydrogenase domain, and a low resolution reconstruction of a construct with the full C-terminus, corroborating that the native oligomeric state of CtBP with bound NADH is tetrameric. Moreover, mutagenesis of key residues involved in interdimer interface diminish the expression of TIAM1 and cell migration in HCT116; CtBP2(−/−) cells supporting a key role of the tetrameric assembly in co-transcriptional function of CtBP2. Thus the tetrameric structure of CtBP2 in solution is validated as the functionally active form of the enzyme.

## Results

### Overall Structure of the minimal dehydrogenase domain of CtBP2, CtBP_31-364_

CtBP2, a 445 residue protein, has required truncations for crystallographic and cryoEM structure determination. The first 30 residues which contain part of the PXDLS motif (Bergman & Blaydes, 2006), are removed from all of our constructs. Crystallization required the removal of the C-terminal (82 residues), which has been shown to be disordered (Nardini et al., 2006). We pursued cryoEM structure determination of CtBP2_31-445_, with the full C-terminus, and CtBP_31-364_ truncation that is equivalent to the construct we crystalized (Hilbert et al., 2014). The CtBP2_31-364_ construct yielded better data with less preferred orientation (Figures S1, S4) and thus is the focus of the detailed analysis. CtBP2_31-364_ was expressed in *E. coli* cells and purified as reported previously (Hilbert et al., 2014; Hilbert et al., 2015). The cryoEM structure determination was performed using both C1 and D2 symmetry, with the D2 symmetry resulting in slightly higher resolution. Reference-free 2D classification reveals distinct classes with different views of the particles (**Figure S1**). The 2D classes also indicate high stability of the tetramer complex. Moreover, reference-free 2D classification only shows classes of the tetramer with no dimeric classes (Figure **S1**). 3D refinement and classification in RELION led to a 3.9Å (FSC = 0.143 criterion) map (**Figure 1**), which improved with per particle CTF refinement to a final resolution of 3.6Å (**Figures S3, S4A-B**). Overall the final EM reconstruction reveals a tetramer of CtBP2 bound with four molecules of NADH (Figure 1C), whose subunit arrangement is very similar to that derived from the crystallographic analysis. Thus, the previously observed tetramer in X-ray crystallographic experiments was not due to crystal contacts, but to the fact that the native state of NADH bound CtBP2 is tetrameric.

**Figure 1.**
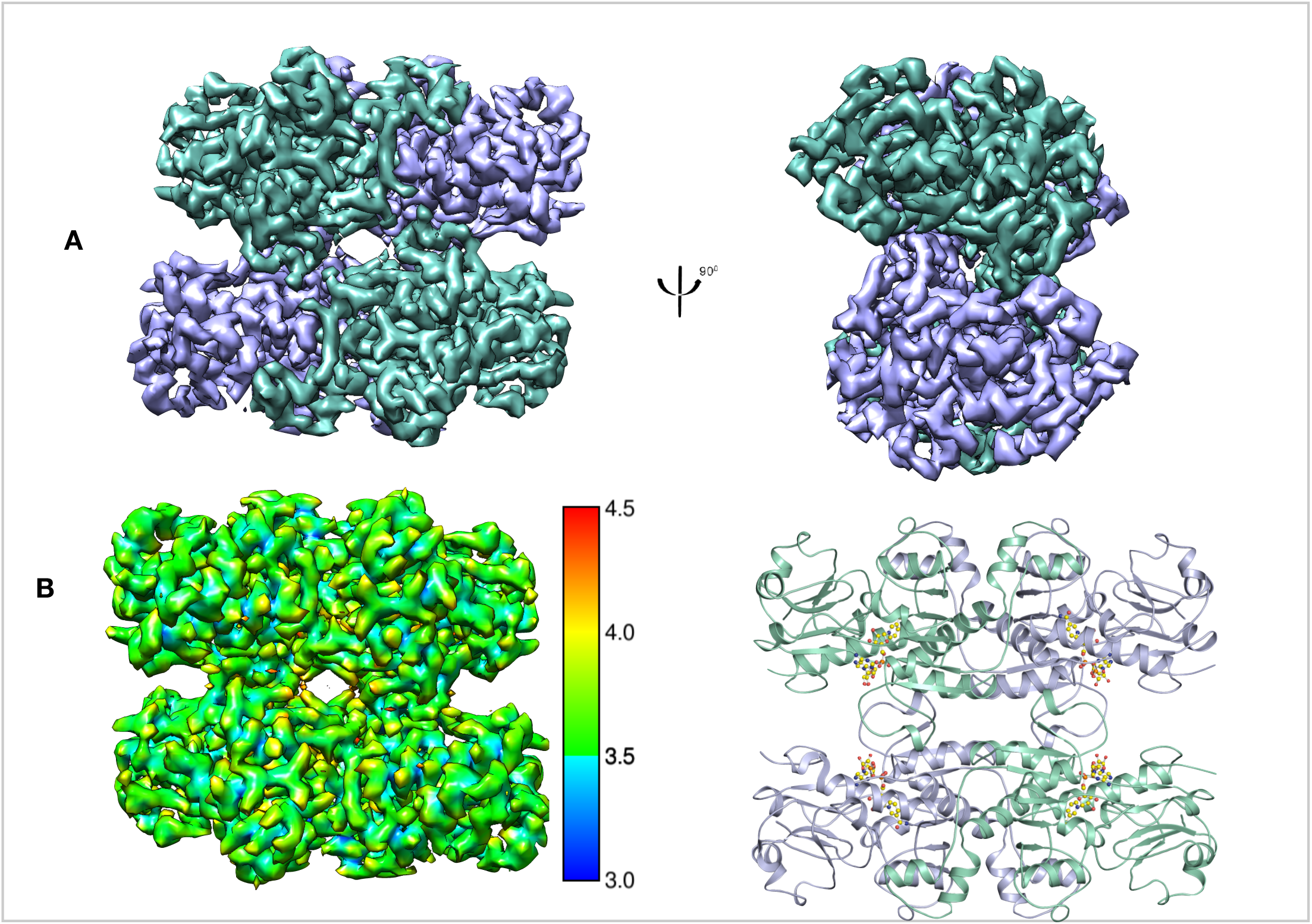
Overall structure of CtBP2_31-364_. (A) Refined and B-factor sharpened 3d reconstruction of CtBP_33-364_ with tetrameric architecture. (B) Local resolution of final map. (C.) Cartoon representation of the EM model with four NADH molecules. Chains A and C are in green cyan, chains B and D are in light blue. **Figure supplement 1A-E:** CryoEM pipeline of CtBP2_31-364_ **Figure supplement 2A-C:** FSC of final reconstruction. Angular distribution of refined particles.

### Overall Structure of CtBP2 with the flexible C-terminal tail, CtBP_31-445_

To tease out the role that the last 80 amino acids of CtBP2 play in tetramer assembly and stability we determined the cryoEM structure of CtBP2_31-445_ (**Figures 2 & S3**). Despite collecting the data under identical conditions as the truncated construct, the longer construct yielded poor quality maps that appear to result from preferred orientation on the EM grids (**Figure S3D**) (Tan et al., 2017). The data was first analyzed without any imposed symmetry hoping to visualize the last 82 amino acids, which are predicted to be highly flexible. Although some 2D and 3D classes showed extra density, this density could not confidently be assigned to the C-terminal flexible tail (Figure **S3C, S4**). Overall, the CtBP2_31-445_ reconstruction performed with D2 symmetry shows tetramers similar to the truncated construct. The first observation made by comparing the 2D classes between **CtBP_31-364_** and **CtBP_31-445_** was that the former had more side views (**Figures S1, S3B**). Consequently, the EM map for **CtBP_31-445_** is limited by preferred orientation of particles (Figure **S3D**). The number of particles in the final reconstruction together with the orientation bias gave rise to a reconstruction with an average resolution of 6Å-12Å. The low quality reconstruction in the presence of the flexible C-terminal domain is likely attributed to a propensity to orient in the EM grids in a small number of preferred orientations.

**Figure 2.**
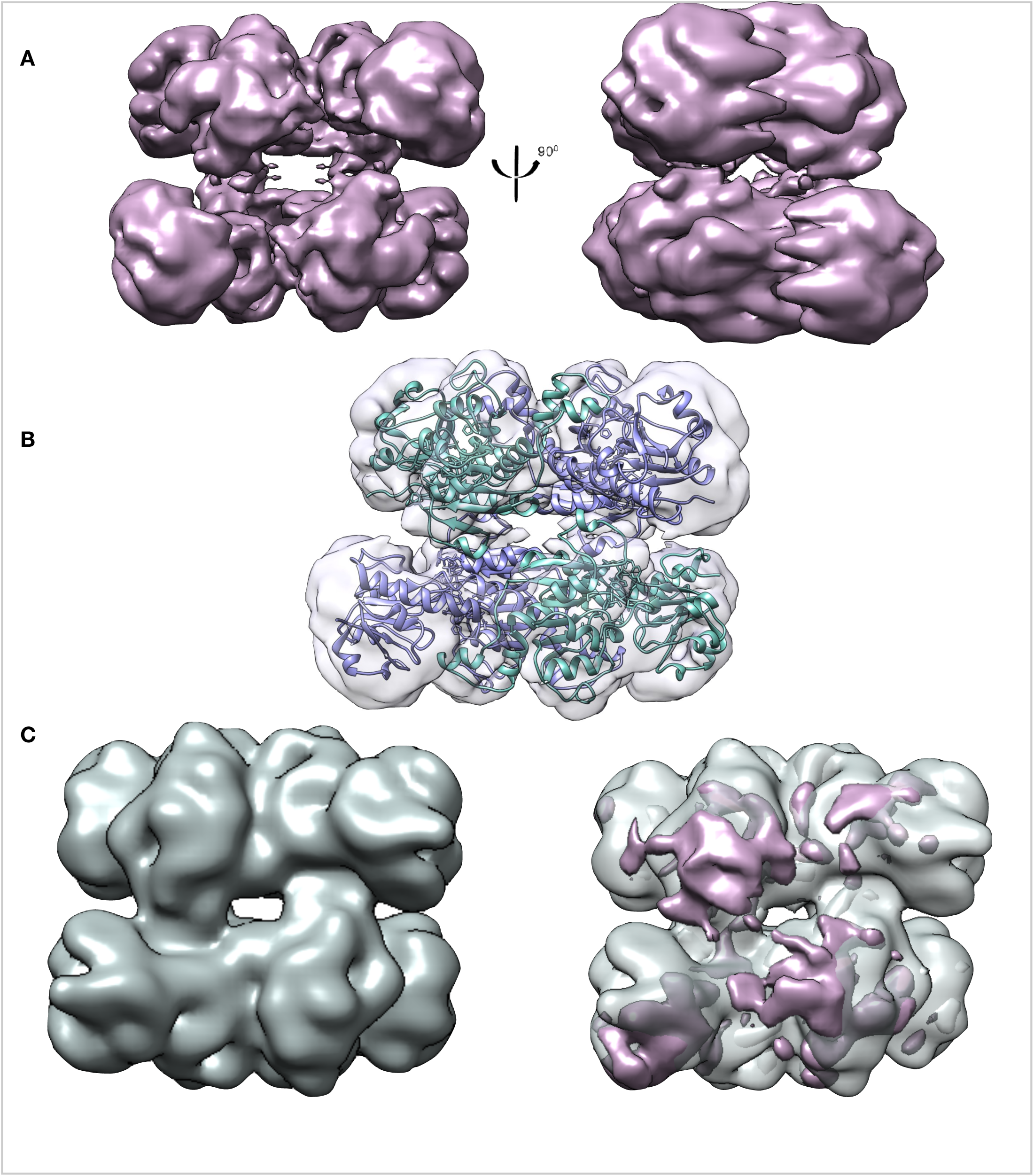
Overall structure of CtBP2_31-445_. (A) Refined and B-factor sharpened 3d reconstruction of CtBP_31-445_ with tetrameric architecture. (B) Fitting of CtBP2_31-364_ EM model into the the CtBP2_31-445_ EM reconstruction. Both the construct with the truncated dehydrogenase domain and the longer construct form stable tetramers. (C) CtBP2_31-364_ map low-pass filtered to 10Å. Diffmap analysis. density (magenta) from CtBP2_31-445_ mapped onto the low-pass filtered map of CtBP_31-364_ **Figure Supplement 3:** CryoEM pipeline of CtBP2_31-445_ **Figure Supplement 4:** CtBP2_31-364_ diffused density

### Comparison of CtBP2_31-364_ to CtBP2_31-445_

To analyze how CtBP2_31-445_ might deviate from the truncated construct a low-pass filtered map of CtBP2_31-364_ (**Figures 2B & S5**) was generated. Difference maps analysis reveals that the monomers in the longer construct are rotated at an angle to bring the NADH biding pocket of each monomer closer together (**Figure 2B**). This observation supports the finding that FL-CtBP proteins formed more stable tetramers as compared to a truncated construct (Madison et al., 2013). Since this cannot be quantified, we hypothesize that rotation of one chain toward the adjacent one would result in a tighter interface, thus a stronger interaction between the two chains. Contrary to previous reports that the last 82 amino acids are required for CtBP2 to assemble into a stable tetramer (Madison et al., 2013) we unambiguously demonstrate that CtBP2_31-364_ forms a tetrameric structure similar to CtBP2_31-445_ (**Figures 1A & 2A**).

### Tetrameric Model of CtBP2_31-364_

The final cryoEM map of CtBP2**_31-364_** provided very clear secondary structural features (**Figure S6**) that permitted rigid-body fitting of the tetrameric crystallographic structure (Hilbert et al., 2014) in Chimera (Pettersen et al., 2004). This fitting was followed by multiple rounds of resulting in a model with high real-space correlation and optimized stereochemical fit (**Table 1**) (**Figure 3**). Pairwise superposition of the two structures yields an average RMSD of 0.8 Å. Cα-Cα distance map analysis (**Figures 3B & S7**) reveals minor differences between the cryo-EM and crystallographic models mainly in loop regions, and interestingly a small rotation in the substrate domain relative to the larger coenzyme domain. Likely due to the stability of the NADH bound complex, comparisons of the two structures reveal that globally these complexes are very similar.

**Figure 3.**
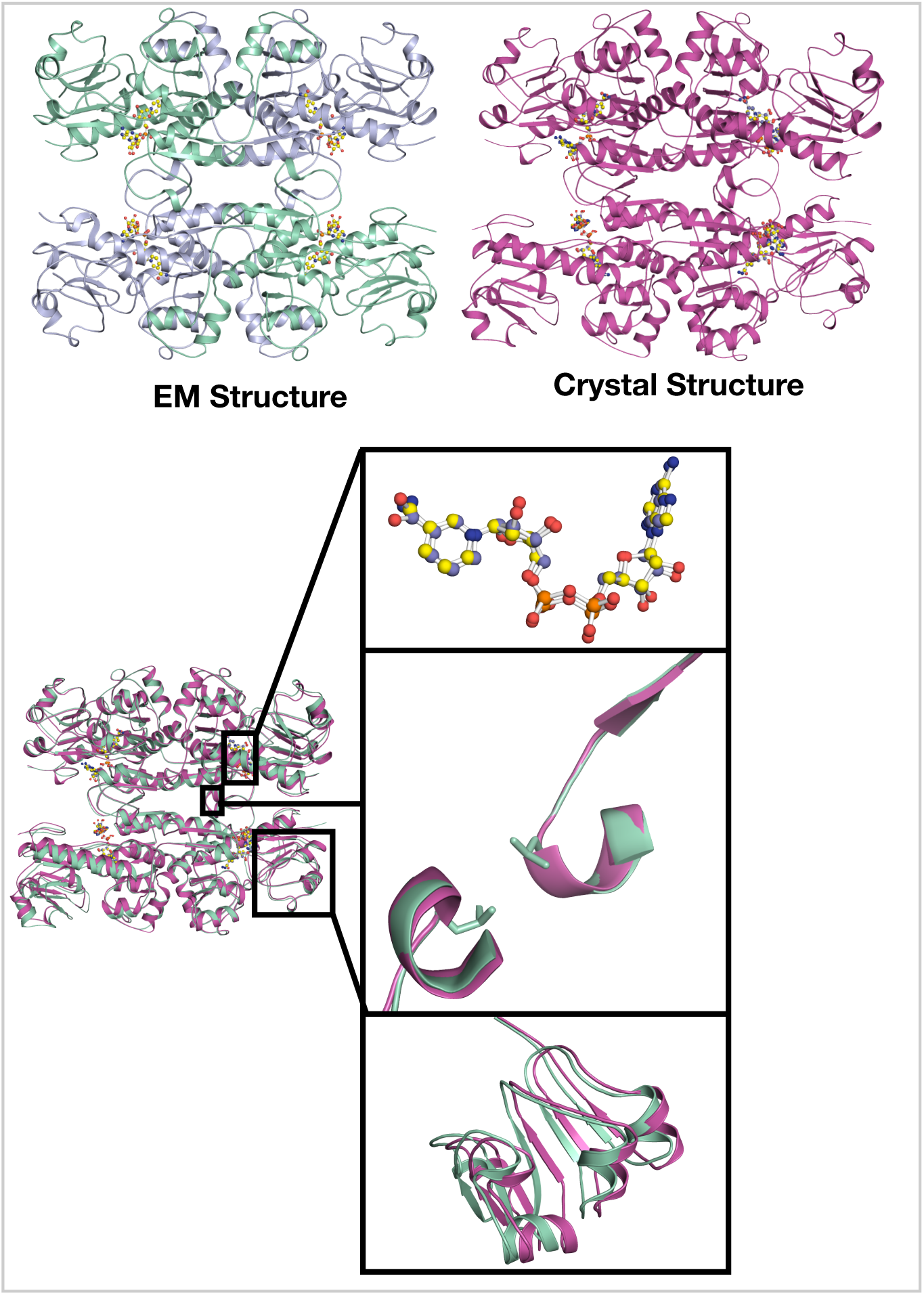
Comparison of the EM and crystal Structures of CtBP2_31-364_. (A) Side by side comparison of the EM model and crystal structures of CtBP2_31-364_. (B) Structural comparison by superimposing amino acids 31-364 of the cryoEM and crystal structures. (C) The NADH molecules of both models align perfectly (D) The interdimer loop with the Leu221 residues superimposes really well onto each other depicting the stability of that region during tetramer assembly. (E) Most of the differences between the two structures lie in the co-factor binding domain. The RMSD between the two structures is 0.781Å. **Figure Supplement6:** Cα-Cα distance map analysis of cryoEM and Crystallography structures.

**Table 1:**
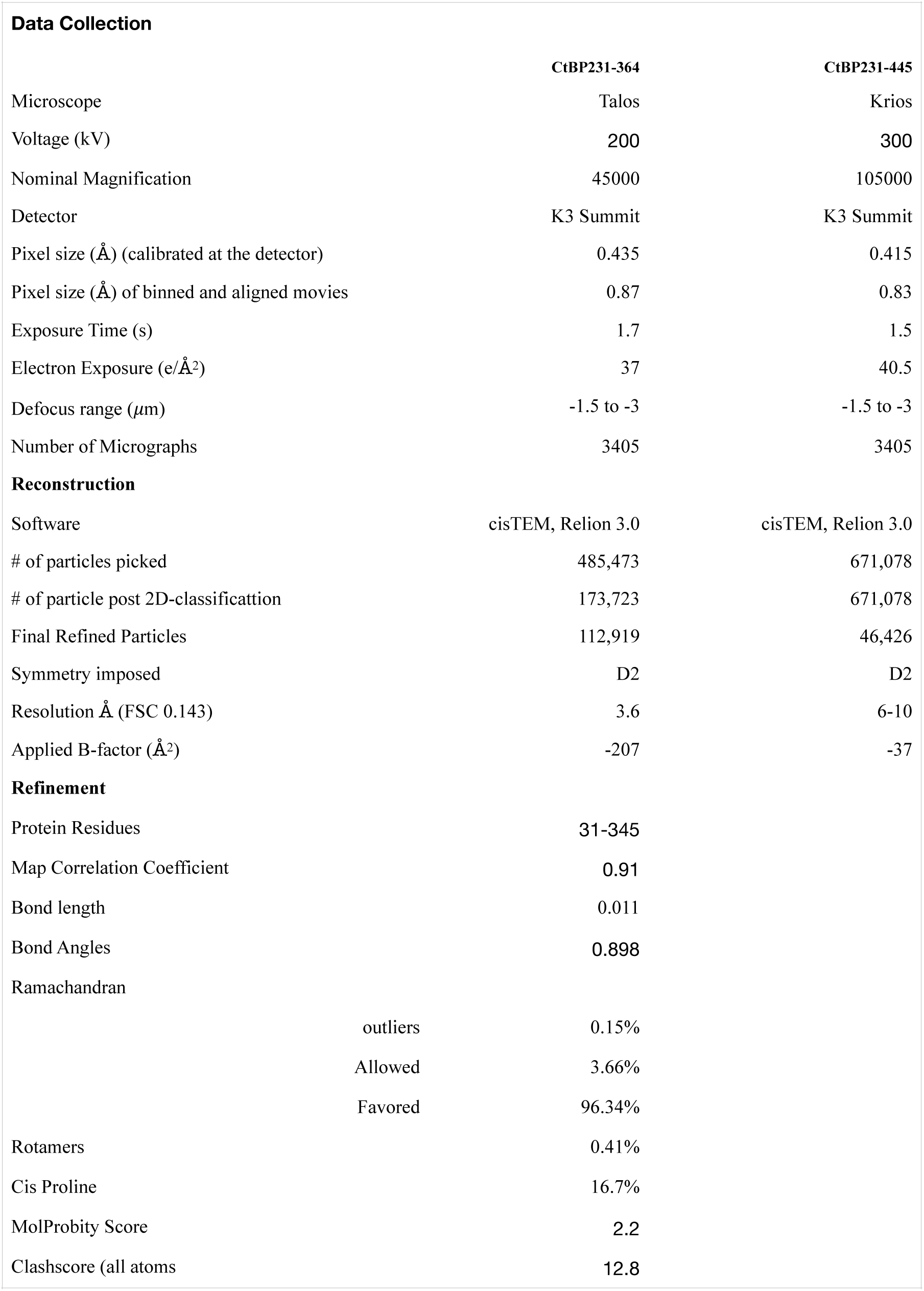
CryoEM data collection, refinement and validation statistics Table.

### Interactions stabilizing CtBP2 tetramer

Most of the structure of CtBP2 was well resolved with the high local resolution of the data set. Density for nearly all the side chains and NADH are clearly visible (**Figure S6**). Each CtBP monomer is composed of a substrate binding domain (31-126, 333-361) and a coenzyme-binding domain (131-325). The overall tetramer with D2 symmetry structure is formed by a dimer of dimers (**Figures 4 & 5**). The most extensive interactions occur at the dimer interface as we reported earlier (Bellesis et al., 2018) and shown here by the PISA analysis in **Table 2** of the cryoEM structure (**Figure 5A & B)**. Each intradimer (AB and CD) buries approximately 3000Å^2^, compared to the 800Å^2^ surface area buried by the interdimers (AD, BC). Based on assembly pathway analyses done on homotetramers, the first complex to assemble is the one with the largest buried surface (J. Chen, Sawyer, & Regan, 2013; Villar et al., 2009) (Quintyn, Yan, & Wysocki, 2015). Consequently, the assembly pathway for NADH-bound CtBP2 is a dimer to a tetramer.

**Figure 4.**
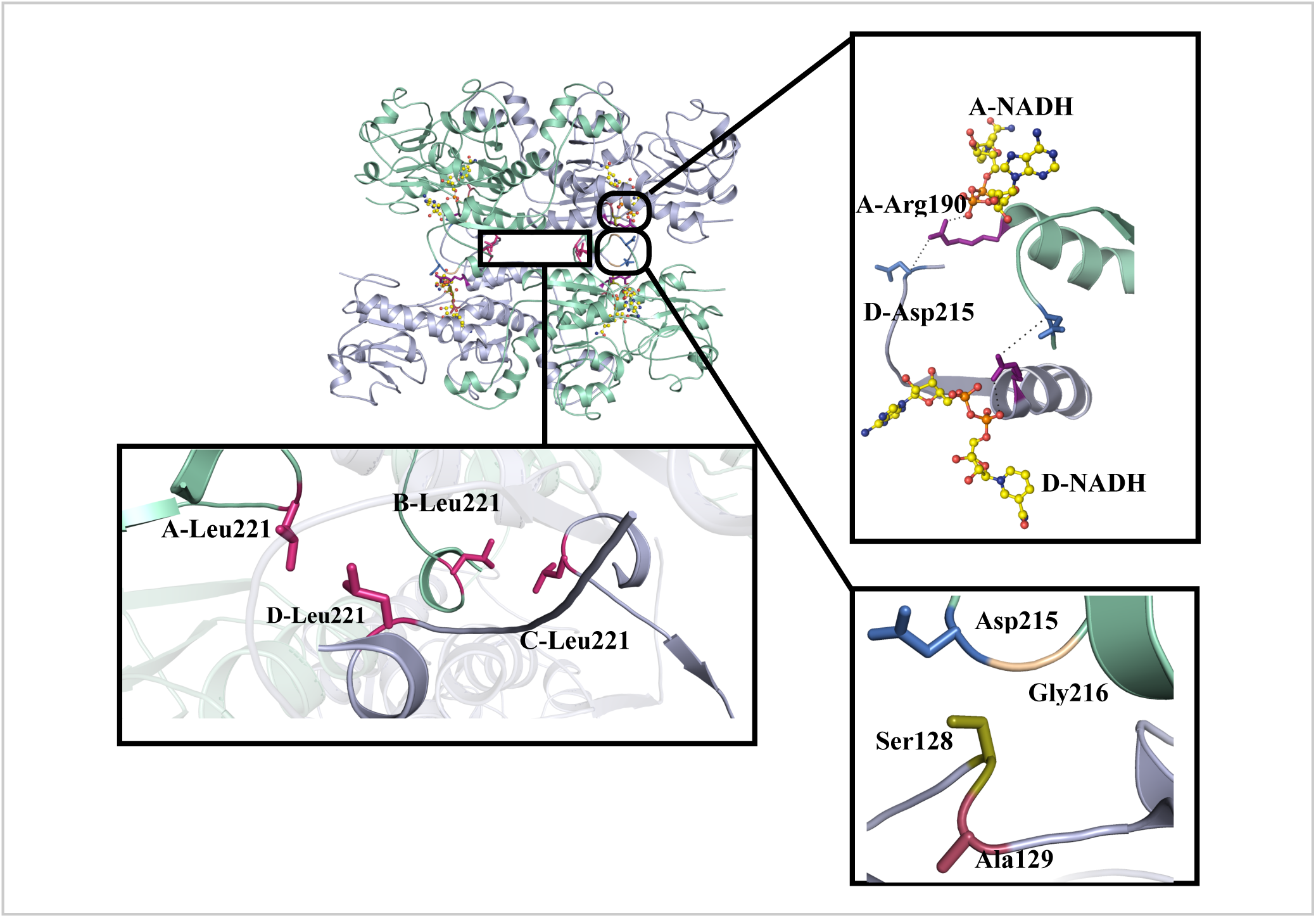
Hydrophilic and hydrophobic interactions that stabilize the tetramer interface. (A) Representative view of CtBP2_31-364_ and the amino acids (B) Zoomed in view of the interactions between NADH (yellow), Arg190 (violet) and Asp215 (marine) across the tetramer interface. (C) Hydrophobic packing of Leu221 stabilizing the tetramer. (D) Representative view of key residues (Ser128, Ala129, GLy216) of the hinge region between the substrate and co-factor binding domains. **Figure Supplement 7:** Representative density of one dimer interface and interactions.

**Figure 5.**
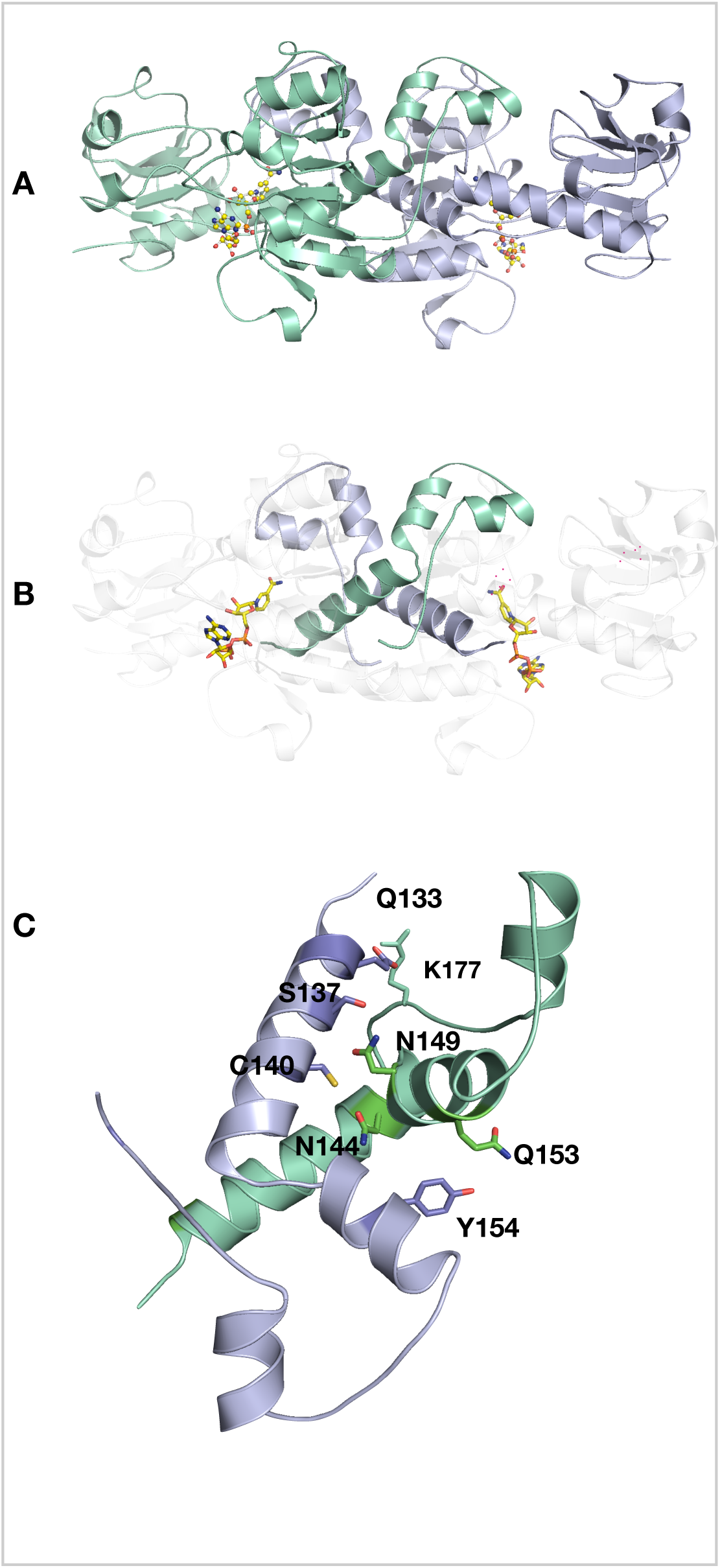
Dimer stabilization. (A) Cartoon representation of the dimer between chain A (light blue) and chain B (greencyan) with two NADH (yellow). (B) For clarity only the dimer interface between the two chains, for clarity only the dimeric interface is shown in green cyan and light blue. (C) Hydrogen-bonding network at the intradimer interface of CtBP_31-364_.

**Table 2:** PISA Analysis of CtBP2_31-364_ CryoEM Structure.

| **Dimer Interfaces** | **Buried area, Å<sup>2</sup>** | **ΔG interface Kcal/mol** | **ΔG Dissociation** | **Salt Bridge** | **H-bonding** | **Hydrophobic** |
| --- | --- | --- | --- | --- | --- | --- |
| AB | 2922 | -41.2 | 36.5 | 17 | 31 | - |
| CD | 3013 | -40.6 | 32.1 | 15 | 39 | - |
| AD | 876 | - | - | - | 9 | 4 |
| CB | 841 | - | - | - | 13 | 7 |

The tetramer interface in the electron density map could be well resolved and side chains placed to explain the tetramerization stability (**Figures 4 & S6**). The tetramer is stabilized by a set of residues**_**clustered near the binding pocket of NADH**_**(Arg190, Leu221), the hinge domains (**Figure 4D**) (Ser128, Ala129) and the hydrophobic clustering of Leu221 at the interdimer interface. Some of these residues either directly contact NADH or interact with each other via a set of hydrogen bonding and hydrophobic interactions (**Figure 4**). For instance, Arg190 participates both in hydrogen bonds with NADH, within its subunit, and the carbonyl oxygen of Asp215’, across the interdimer interface (**Figures 4B & S6B**). Another set of interactions that stabilize the tetramer is the hydrophobic packing of the side chains of Leu221 (**Figure 4C**) located at the interdimer interface. Mutations of these residues that disrupt the tetrameric assembly (Bellesis et al., 2018) provide an opportunity to investigate that role of tetramer stability and CtBP transcriptional activity.

### Tetramer Destabilizing Mutants Decrease Cell Migration

Our cryoEM results establish that the tetrameric form of CtBP2 deduced from the crystal structure does represent the solution tetrameric structure and is not an artifact of the CtBP2 crystal lattice. Given the evidence correlating NADH binding and oligomerization with activation (Madison et al., 2013) (Bellesis et al., 2018), the stable tetrameric form of CtBP2 identified here is likely to be functionally important. To directly test the hypothesis that our observed tetrameric form is the functionally active form, plasmids encoding a series of CtBP2 mutants that have previously been shown to inhibit tetramer formation (Bellesis et al., 2018) were transfected into HCT116 colon cancer cells that have had CRISPR-mediated deletion of both CtBP2 alleles (Chawla et al., 2018). These transfected cell populations were then investigated for expression of the CtBP2 transcriptional target gene TIAM1 and induction of cell migratory activity, two cellular functions specifically correlated with oncogenic activity (Dcona et al., 2017; Ma et al., 2020).

TIAM1 promotes cancer progression and metastasis, and CtBP2 is a key driver of TIAM1 transcription (Paliwal et al., 2012). To investigate the role of the CtBP2 tetrameric assembly in transcriptional activation of TIAM1, plasmids encoding CtBP2_WT_, empty vector and tetramer destabilizing mutants (S128T, A129L, R190Q, G216N, L221Y) (**Figure 4**) were transfected into HCT116; CtBP2(−/−) cells. Monitoring of transfected cell lysates by CtBP2 immunoblot indicated equivalent expression of CtBP2_WT_ and all five mutant proteins, as determined by densitometry of the CtBP2 and GAPDH loading control immunoblots (**Figures S8 & S9**). Following transfection, total RNA was extracted and TIAM1 mRNA abundance was determined by qPCR, as described in the Experimental Procedures. As shown in **Figure 6A**, transfection with pCtBP2_WT_ resulted in more than a 2-fold increase of TIAM1 mRNA expression over the empty vector control. In sharp contrast, all five tetramer destabilizing mutants were defective for TIAM1 induction compared to CtBP2_WT_ (p<0.01 for comparison of TIAM induction by CtBP2_WT_ vs. each mutant). Indeed, each of the mutants appeared to repress TIAM1 transcription below basal levels seen with vector transfection, but the comparison of TIAM1 expression between vector and mutant CtBP2 transfection did not achieve statistical significance. Thus, CtBP2 mutants that are incapable of forming tetramers are also deficient in the transcriptional activation of TIAM1.

**Figure 6.**
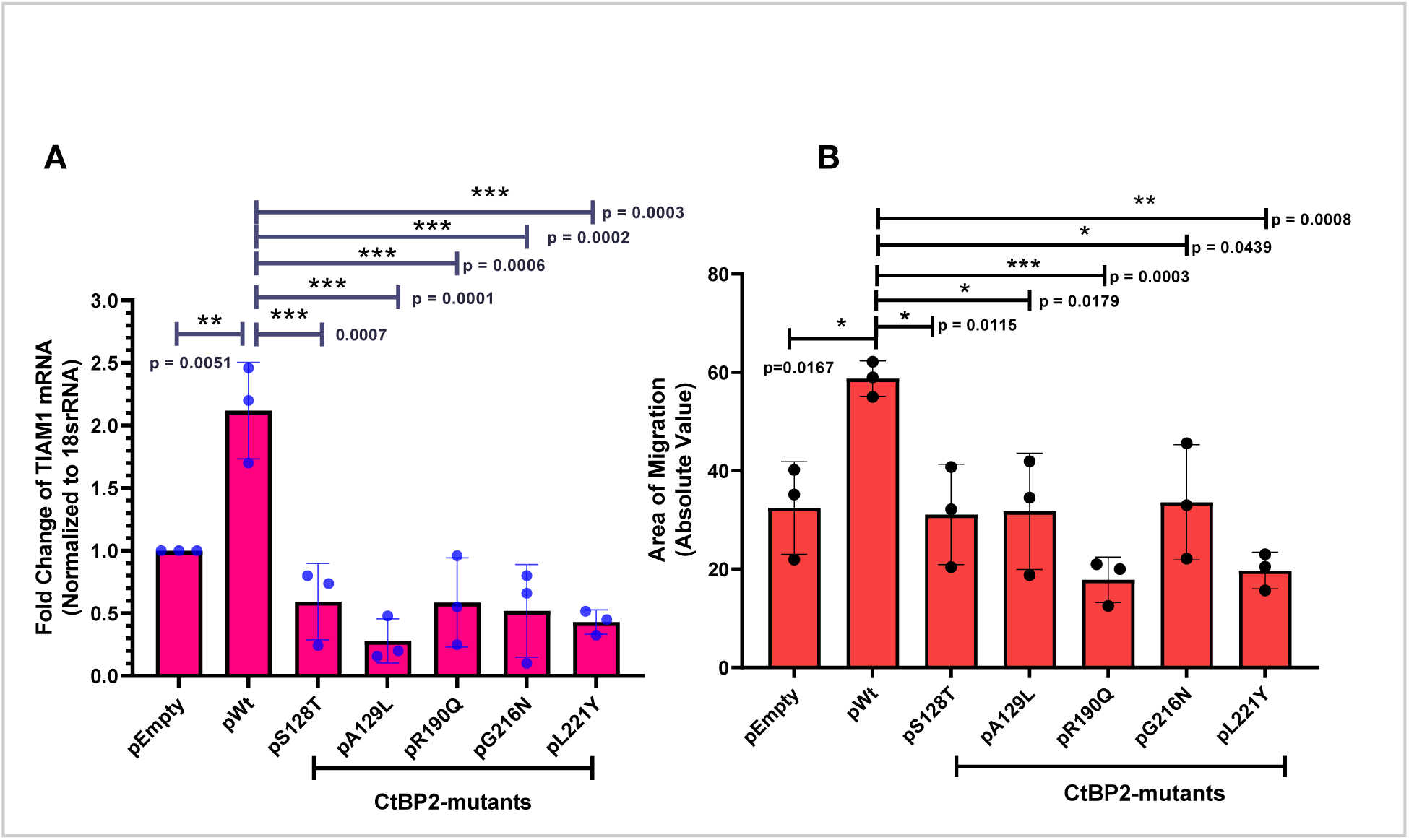
Tetramerization of CtBP2 is critical for co-transcriptional activity. HCT116 CtBP2(−/−) cells were transfected with either plasmids encoding either WT CtBP2_WT_ or tetramer destabilizing mutants (S128T, A129L, R190Q, G216N and L221Y). (A) RT-qPCR analysis of TIAM1 gene expression. Fold change of expression was normalized to 18sRNA. (B) Quantification of area of migration after scratch assay. All assays were repeated N=3 times, and statistical significance was calculated using one-way ANOVA, where * indicates p ≤ 0. 05; ** p ≤ 0.01; *** p ≤ 0.001.Asterisks indicate significant difference between the different variants. **Figure Supplement 8:** Expression level of CtBP2 variants.

Cellular migration is an important hallmark of oncogenesis and CtBP2 robustly induces cell migration and invasion in cell culture as a correlate of CtBP’s in vivo activities in promoting invasion and metastasis (Dcona et al., 2017). To test the contribution of the CtBP tetramer to the CtBP induction of cellular migration, a “scratch assay” was used in which HCT116; CtBP2(−/−) cells transfected with the vectors described above were grown to confluence and then a scratch was made on the plate, and cells were allowed to migrate into the scratched area, and closure of the scratch due to cell migration was quantified after 24 hours. As shown in **Figure 6B**, closure of the scratch increased from a basal value of 40% with empty vector transfection to approximately 60% with the transfection of CtBP2_WT_ expression vector. All five tetramer destabilizing mutants exhibited defective induction of migration compared to CtBP2_WT_ (p<0.05). As for TIAM1 transcription, certain of the mutants appeared to exert a dominant negative effect, driving migration below basal levels, but these differences did not achieve statistical significance. These results, thus, provide the first direct evidence that the tetrameric form of CtBP2, observed in the cryoEM reconstruction presented here, is required for co-transcriptional activity regulating TIAM1 expression and induction of cell migratory behavior.

## Discussion

Eukaryotic gene transcription is regulated on many levels. One form of regulation involves recruitment of transcriptional factors and regulators that assemble into macromolecular complexes (Soutourina, 2018; Vilar & Saiz, 2005). Our structural analyses on CtBP2_31-364_ and CtBP2_31-445_ reveal a tetrameric assembly for the CtBP2 protein. The observed tetramer is assembled from two dimers, each stabilized by both hydrophilic and hydrophobic interactions with an extensive interaction area of over 3000Å^**2**^. In contrast, the tetramer formation is mediated by interacting loops from adjacent dimers with significantly lower interaction area, 800Å^**2**^. The model corroborates the previous crystallographic model that predicted the tetramer as the oligomeric state for both CtBP1 and CtBP2 (Bellesis et al., 2018). Our Cα-Cα distance analysis reinforces that the EM structure is very similar to the crystal model with no significant domain reorientation or movement due to the protein being in solution. As observed by Nardini et al, NADH binding locks t-CTBP/BARS into a closed conformation in which the substrate and co-enzyme binding domains are in close proximity (Nardini et al., 2003).

Previous studies suggested that although CtBP2_31-364_ can form tetramers, the dimeric species would be more prevalent because the last eighty residues are missing. In our cryoEM analysis, the only presented classes are tetrameric with no dimers observed. One major reason for that is the final concentration of the cryoEM sample will be higher compared to the concentration used for MALS and other biochemical assays, which will result in a higher binding between the subunits.

Despite the difference in resolution, the overall EM reconstruction for both CtBP2 constructs was highly similar. The main difference lies in the orientation of the coenzyme binding domain of CtBP2_31-445_, which has a slight rotation toward the adjacent chain. This rotation brings the two chains closer together and will consequently lead to tighter interactions. This result suggests that previous results showing greater tetramer formation in the presence of the C-terminal residues (Madison et al., 2013) may not result from specific intersubunit interactions involving the C-terminus, but rather from an effect of the disordered C-terminal domain altering domain orientation.

Our cryoEM studies not only confirm the tetrameric assembly, but also show that CtBP2 bound to NADH forms a stable complex. The latter is important because of CtBP’s role in promoting the assembly of higher order complexes such as the CtBP-mediated repression complex (Turner & Crossley, 2001) (Good, Zalatan, & Lim, 2011; Sun & Fang, 2016). As the hub for assembly of this complex, the tetramer stability and rigidity may be essential for the assembly of other co-factors. Future studies of fulllength CtBP2 in complex with interacting partners may provide a detailed structural understanding of the role of the C-terminal domain both in tetramer stability and assembly of CtBP-mediated transcriptional complexes.

CtBP2 proteins have been shown to directly activate *Tiam1* to facilitate EMT transition and cancer progression (Di et al., 2013; Paliwal et al., 2012). Our cell-based assay analysis of tetramer destabilizing CtBP2 mutants demonstrates that tetrameric assembly is required for transcriptional activity. As shown in **Figure 6**, tetramer-destabilizing mutants abrogate induction of TIAM1 expression (**Figure 6A**), consequently affecting the ability of those mutants to induce cell migration (**Figure 6B**). The observation that tetramer-destabilizing mutants exhibit a possible dominant negative affect by lowering TIAM1 expression or cell migration below that seen with empty vector control is intriguing but the effect did not achieve statistical significance in our experiments. While beyond the scope of the present work, we speculate that such a dominant negative effect could result from CtBP2 forming dimers with endogenous CtBP1 that are then unable to assemble into tetramers.

The co-transcriptional activators CtBP1 and CtBP2 have been extensively studied because of their implication in various cancers. High expression of CtBPs in cancerous cells is linked to poor outcomes (Bergman & Blaydes, 2006). Delineating the relevant assembly of these proteins in the cell is of critical importance for the development of targeted therapies that act through disruption of the CtBP tetramer.

## Experimental Procedures

### Expression of CtBP2_31-364_ and CtBP2_31-445_

The expression and purification procedures were adapted and optimized from earlier studies (Hilbert et al., 2014). The ligated, purified plasmid containing the desired CtBP construct was transformed into Z-competent BL21(DE3) RIL E. *coli* cells. A single clonal colony was then grown in a starter culture of LB broth overnight at 37C. The starter culture was used to inoculate between three and six 1L cultures grown in Research Products International Terrific Broth using 50mL starter per liter. Cultures were grown at 37C while shaking at 150RPM and induced with 1 mL 0.2 M IPTG after reaching OD_600_ between 0.800 and 1.00. The temperature was reduced to 30C at the time of induction and the cells were harvested four hours later. The cells were pelleted by centrifuging for 20 minutes at 4700 RPM, and resuspended in 10 mL harvesting buffer (pH 7.6; 0.1 M NaCl; 0.05 M Tris-HCl; 0.2 mM EDTA) per liter of culture. One tablet of EDTA-free complete Mini (Roche Diagnostics) protease inhibitor cocktail was added per liter of culture.

### Purification of CtBP2_31-364_ and CtBP2_31-445_

Cells were thawed slowly on ice and then lysed in a Microfluidics Corporation model 1109 cell disrupter. 35 mg of Roche Diagnostics DNase I, 500μL 2M MgCl2 and 500μL 40 mM CaCl2 were added per 100mL lysate. The lysate was then gently stirred at 4C for 30 minutes. The insoluble fraction was pelleted at 19,000RPM for 45 minutes. The supernatant was then mixed with 8 mL HisPur^TM^ Ni-NTA Resin (Thermo Scientific), and gently stirred at 4 for two hours to allow CtBP to bind to the resin.

The bead-supernatant mixture was placed in a BioRad Econo-Column® at 4C and the soluble fraction was allowed to flow through. The beads were then cleaned with 40 mL wash buffer (0.0625 M Tris:HCl pH 7.4; 0.375 M NaCl; 0.05 M imidazole; 0.625 mM EDTA; 1.0 mM DTT), followed by 50 mL of wash buffer supplemented with an additional 1.7 M NaCl. Another 10 mL wash buffer was passed over the beads before 50 mL wash buffer supplemented with 0.5% Triton-X 100 was added. An additional 10 mL of wash buffer again followed. CtBP was eluted from the beads using 25 mL wash buffer supplemented with 250 mM imidazole. The protein was then concentrated by centrifuging at 5000 RPM in an Amicon® Ultra-15 10K centrifugation column (Millipore). Protein concentration was measured by UV absorbance at 280nm using an Ultraspec 2100 pro by Amersham Biosciences. The protein sample was further purified by FPLC. The FPLC (ÄTKAprime plus by GE Healthcare) and size exclusion column (Highload™ 16/60 Superdex™ 200 prep grade) were equilibrated with “FPLC Buffer” (50mM Tris:HCl pH 7.4, 300 mM NaCl, 5 mM EDTA, 2 mM DTT). The sample was prepared by adding 1.5 mM NADH to the concentrated protein solution

The solution was then centrifuged at 8000RPM for six minutes at 4 to remove any small insoluble fraction. The flow rate was set to 1 mL/min and 62 fractions of 2 mL each were collected the appropriate fractions were concentrated in an Amicon® Ultra-15 10K centrifugation column.

### Sample preparation of CtBP2_31-364_ and CtBP2_31-445_ for CryoEM

For our cryoEM studies, two microliters of purified sample at a concentration of 250nM CtBP2_31-364_ and 500nM CtBP2_31-445_ was added to glow-discharged, 200-mesh C-flat 1.2/1.3 EM grids. The sample was blotted for 4s at 4°C under 95% humidity and vitrified in liquid ethane cooled by liquid nitrogen using the Vitroblot Mark IV.

### Image Acquisition

The dataset for CtBP2_31-364_ were recorded on the Talos Arctica operated at 200 kV equipped with a Gatan K3 summit direct electron detector operating in electron counting mode with −1.5 to −3**μ**m defocus. Automated data acquisition was carried out using SerialEM (Mastronarde, 2005) at a nominal magnification of 57,000X with a pixel size of 0.435A or CtBP2_31-364_. In Total 3405 movies were recorded for CtBP2_31-364_. 29 frames of movies were collected with a defocus range of −1.3 to −1.5 **μ**m at a total dose of 37 electrons per Å^2^

The images for CtBP2_31-445_ were acquired on the Titan Krios (300kV) equipped with a Gatan K3 summit direct electron detector operating in electron counting mode with −1.5 to −3**μ**m defocus. As above, automated data acquisition was carried out using SerialEM at a nominal magnification of 105,000X and a pixel size of 0.415A, respectively. 4752 movies were collected for CtBP2_31-445_ with a total of 25 frames at a total dose of 40 electrons per Å^2^

### Data Processing

Super-resolution movie frames were binned to the physical pixel of 0.87 and 0.83 for **CtBP2_31-364_** and **CtBP2_31-445_**, respectively. Alignment and beam-induced motion correction was done using IMOD (Kremer, Mastronarde, & McIntosh, 1996). Contrast transfer function (CTF) parameters were estimated with CTFFIND4, reference-free particle picking was conducted in the cisTEM software package (Grant, Rohou, & Grigorieff, 2018; Grigorieff, 2016). For **CtBP2_31-364_** 485,473 particles were selected from 3405 micrographs and extracted with a box of 256 pixels. The 671,078 particles of **CtBP2_31-445_** from 4752 micrographs were extracted with a box of 300 pixels. For both datasets, reference-free 2D classification with no imposed symmetry was carried out to attest the quality and homogeneity of the data. Three rounds of 2D classification were performed to further purify the CtBP2_31-445_ data set.

### 3D reconstruction of CtBP2_31-364_

Selected 2D classes were pooled to generate an *Ab initio* map in C1 in cisTEM. The particle stacks, CTF parameters and ab-inition models were transferred to RELION 3.0.2 for further processing (Zivanov et al., 2018). 3D classification yielded three classes (Sup. Fig. 3), the best class was extracted and further refined to higher resolution. All 3D classification and refinement were repeated in D2. A soft binary mask was generated and used for another round of 3D classification and refinement. Additionally we used per particle CTF estimation as implemented in RELION 3.0.2 to generate the 3.6Å map. B-factor determination was done automatically in RELION. Reported resolutions are based on the gold-standard Fourier Shell Correlation (FSC) 0.143 criterion. Data processing and analysis for CtBP2_31-445_ were performed similarly to the shorter construct. Due to the high noise level of the post-processed map, possibly due to over-masking, we tested different B-factor values to arrive at a decent reconstruction. The final reconstruction had an applied B-factor of −37Å^2^. Difference map analysis between CtBP2_31-445_ an d CtBP2_31-364_ was done using diffmap (http://grigoriefflab.janelia.org/diffmap)

### Model Building and Refinement

Rigid body fitting was carried out in Chimera (Pettersen et al., 2004) by docking the CtBP2 tetramer model (PDB ID: 46UQ) into the map density. As a starting model we used the crystallographic tetramer for refinement. First round of refinement and model building were done in COOT **(V0.8.9.2) (Emsley & Cowtan, 2004)**. The model was further refined using Phenix.real_space_refine and stereochemistry was validated using phenix.molprobity (Table 1) (Adams et al., 2010) (V. B. Chen et al., 2010). Figures were generated in pymol and chimera. Resmap was used for local resolution estimation (Kucukelbir, Sigworth, & Tagare, 2014). To further parse out differences between the two models, we performed distance map analysis by comparing Cα-Cα distance between every pair of amino acids of the EM and crystal structures using Chimera RRdistMaps (J. E. Chen, Huang, & Ferrin, 2015). Interface and assembly analyses were calculated by PISA analysis (Krissinel & Henrick, 2007). The EM model was uploaded and analyzed on the PDBe PISA (v1.52) analysis software.

### Antibodies and Immunoblot Analyses

Antibodies used in immunoblotting (IB) assay are CtBP2 antibody (discontinued by SCBT; E-16: sc-5966) and GAPDH antibody (SCBT, sc-47724). Antibodies were used at dilutions suggested by the manufacturers. For immunoblot analysis, 25 μg of total protein extract was boiled at 95°C in sample buffer, followed by separation on SDS-PAGE (Novex gels, 4-12% Bis-Tris), and then transferred onto nitrocellulose membrane (0.45-mm porosity) (GE Healthcare). The membrane was incubated for 1 to 2 hours in blocking buffer [Tris-buffered saline, 0.1% Tween (TBS-T), 5% nonfat dry milk], followed by incubation overnight at 4°C with the primary antibody solubilized in blocking buffer with sodium azide (0.01%). After 3X washes of 5 minutes with TBS-T, the blot was incubated with Alexa Fluor 680 or 790 nm secondary antibodies (Invitrogen) for 1 hour in TBS-T and visualized on a Bio-Rad imager.

### Cell Culture and Transfection

HCT116; CtBP2(−/−) (Chawla *et al.,* 2018; Dcona *et al.,* 2019) were maintained in Dulbecco’s modified Eagle’s medium (DMEM) supplemented with 10% (v/v) FBS and penicillin–streptomycin in a humidified incubator equilibrated with 5% CO2 at 37°C. Cells were authenticated by examination of morphology and growth characteristics and were confirmed to be mycoplasma-free using DAPI-staining and PCR. Transfections were performed using a standard protocol for Lipofectamine-2000 (LP2000) (Thermo-Fisher) based plasmid delivery. Briefly, required concentrations of plasmids and LP2000 were solubilized in 100 μL of Opti-MEM media in different tubes. After 10 minutes of incubation at room temperature, the contents from tube containing LP2000 were pipetted into the tube containing plasmid solution to form a complex. After further incubation of 30 minutes, the complex was pipetted into the media to transfect HCT116; CtBP2(−/−) cells.

### Site-directed Mutagenesis and PCR

The CtBP2_WT_ mammalian expression vector expresses full-length CtBP2 in a modified pcDNA3.0 mammalian expression vector that also expresses GFP and puromycin acetyltransferase (PAC) as a chimeric protein (M. Michael Dcona et al., 2019). A QuikChange site-directed mutagenesis protocol was used to generate the tetramer-deficient CtBP2 mutants using the CtBP2_WT_ expression vector as a template, namely, S128T, A129L, R190Q, G216N and L221Y.

### RT-PCR and real-time RT-PCR

48 hours post-transfection with 6-8 μg of empty-vector, CtBP2_WT_, S128T, A129L, R190Q, G216N or L221Y plasmids into HCT116; CtBP2(−/−) cells, total cellular RNAs were isolated from samples using QIAGEN RNeasy kit and instructions therein. Later, cDNA synthesis was carried out using SensiFAST cDNA Synthesis Kit from BIOLINE. Quantitation of all gene transcripts was done by qPCR using SYBR Green (Applied Biosystems, Foster City, CA) and an ABI 7300 (Applied Biosystems) machine. 18srRNA expression was used as an internal control.

The primer pairs used were:

18srRNA: 5’-CGCCGCTAGAGGTGAAATTC-3’ (forward) and
5’-TGGCAAATGCTTTCGCTCTG-3’ (reverse);
TIAM1: 5’-CGCTGGAGTCGTACCTCATC-3’ (forward) and
5’-GGTCAAACACAGCCCCAAAC-3’ (reverse)

Relative amounts of the mRNA transcripts were calculated using the ΔΔCT method and reported as fold change with respect to empty-vector transfection(Schmittgen & Livak, 2008). The experiments were repeated N=3 times and the statistical significance was calculated using one-way ANOVA.

### In Vitro Wound-Healing Assay

HCT116; CtBP2(−/−) cells were seeded into six-well dishes at a density of 5 × 10^5^ cells/well. The dishes were cultured as confluent monolayers and were then transfected with 3 μg of EV, CtBP2_WT_, S128T, A129L, R190Q, G216N or L221Y expression plasmids. 24 hours post-transfection, the equivalent transfection efficiency of all plasmids was confirmed by indirect fluorescence microscopy for GFP, and a scratch was then made once per well with a 200-μl pipette tip to create an artificial wound. wounded cell cultures were then incubated in the presence of DMEM after thorough, but gentle washes. The migration of cells was monitored over a duration of 24 hours as a function of how far from the scratch line the cells had progressed. The scratch closures were quantified using ImageJ (NIH) using wound-healing macros. The area at a time point is normalized relative to 0-hour time and reported as absolute value. The experiments were repeated N=3 times and the statistical significance was calculated using one-way ANOVA.

## Acknowledgements

We are grateful to Dr. Gabriel Demo for his mentorship in teaching AMJ single particle cryoEM analysis. His teaching has been invaluable to AMJ’s ability to conduct the analysis for this paper. We thank Dr. Anna Loveland for valuable discussions and feedback on the manuscript. We thank members of the Kelch Lab: Dr. Christl Gaubitz, Dr. Nicholas Stone and Dr. Janelle Hayes for insightful discussions on data analysis and sharing resources. We thank Drs. Kyounghwan Lee, KangKang, Song and Chen Xu for their help with data collection at the UMMS cryoEM Facility. Many thanks to Mr. Florian Leidner for setting up our workstation. We thank Dr. Benjamin Morris for creating the A129L mutant mammalian expression vector used in the cell-based assays. Services in support of the research project were generated by the VCU Massey Cancer Center Biostatistics Shared Resource, supported, in part, with funding from NIH-NCI Cancer Center Support Grant P30 CA016059. This work is supported by NIH grant R01 GM119014 to WER and NIH grant F31 GM129988 to AMJ.

## Data Availability

CryoEM Maps for CtBP2_31-364_ and CtBP_31-445_ were deposited to the EMDB with accession codes EMD-21811 and EMD-11015, respectively. The Atomic coordinates for CtBP2_31-364_ have been deposited in the PDB under ID 6WKW.

