## Supplemental Figures for "CryoEM Structure of CtBP2 Confirms Tetrameric Architecture"

**Figure S1A**

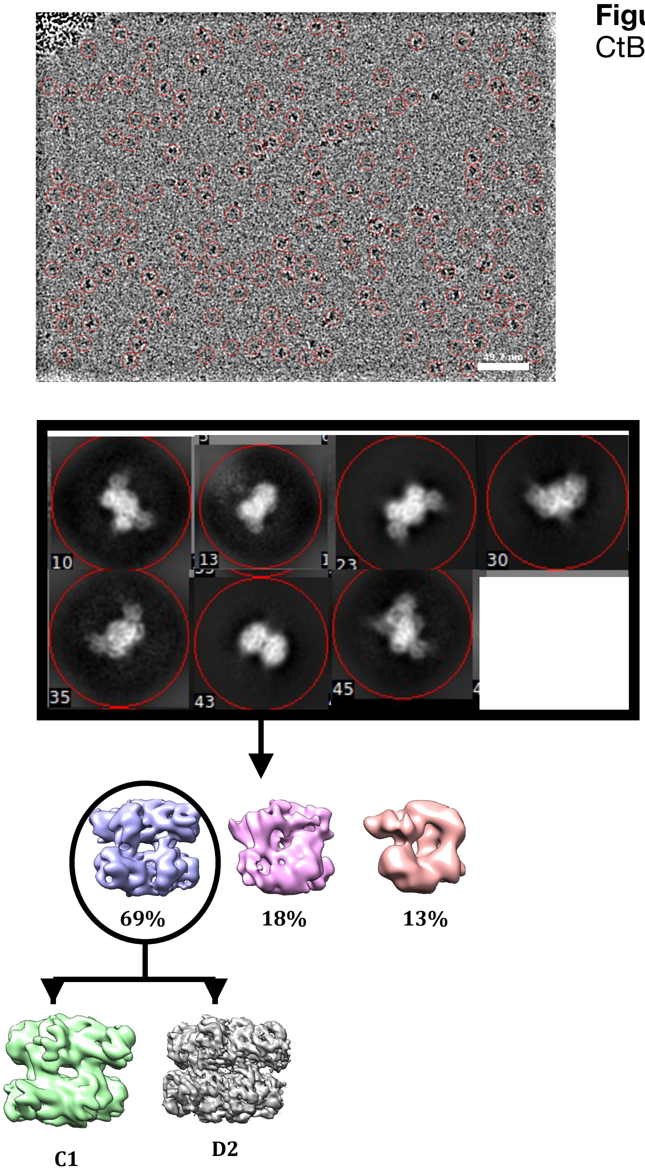


**Figure S1A-E**: CryoEM pipeline of CtBP2$_{31\text{-}364}$

#### Figure S1C

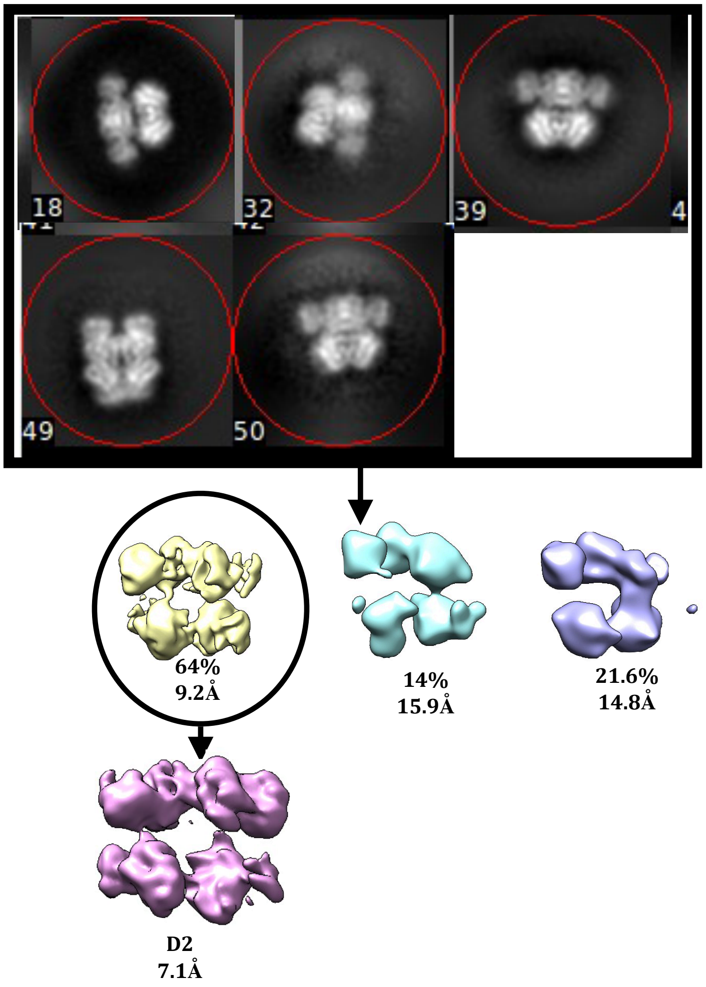


#### Figure S1D

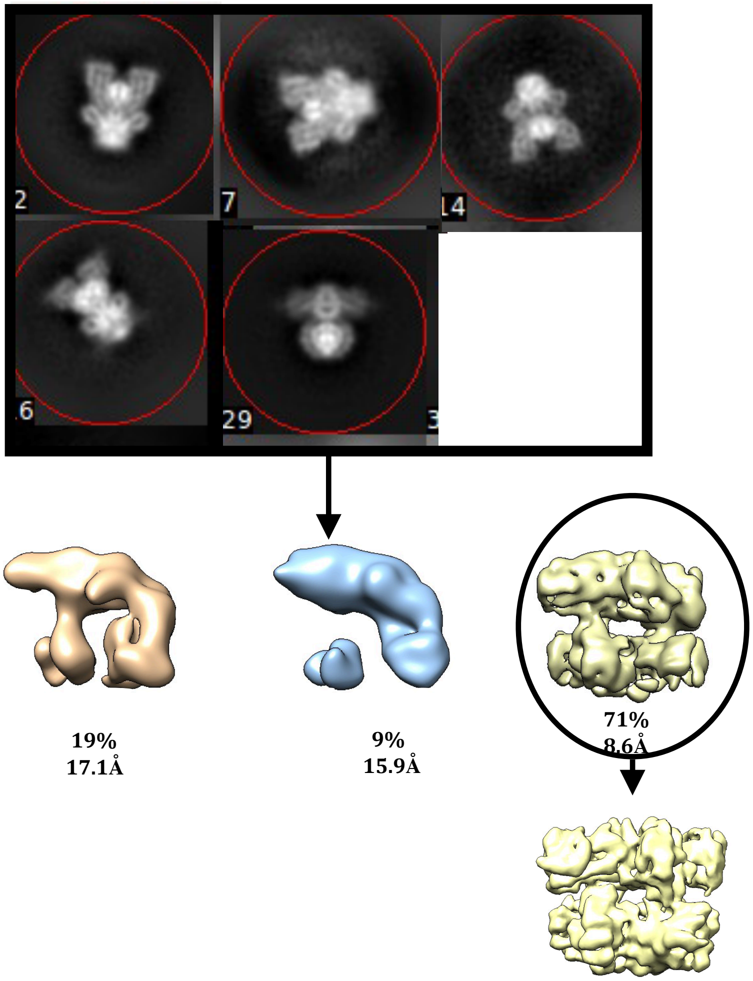


#### Figure S1E

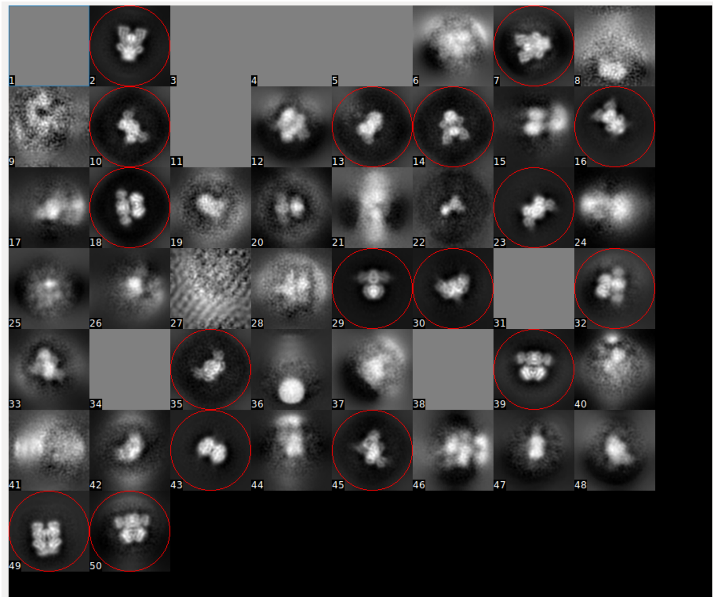

### Figure S1F

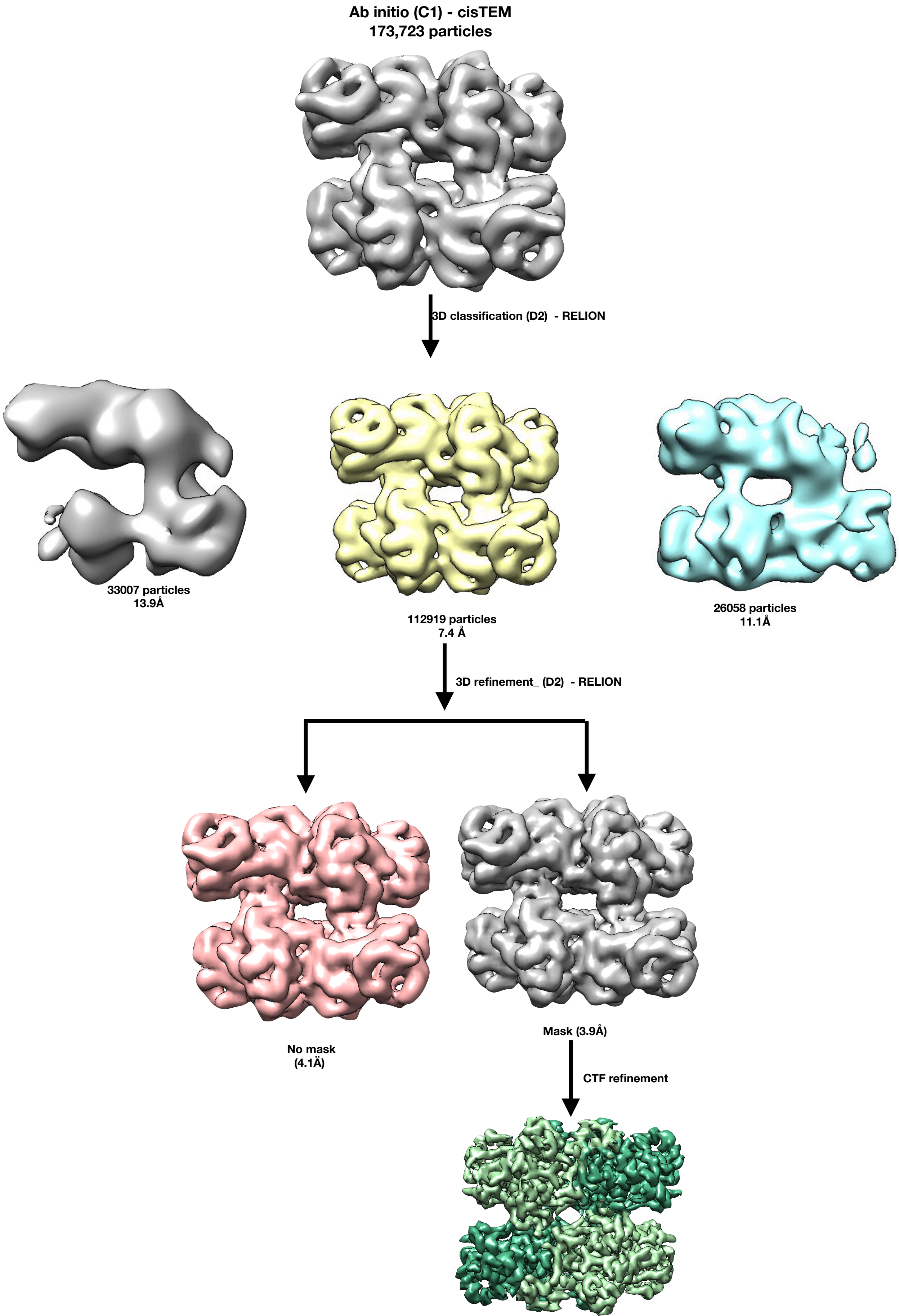
Ab initio (C1) - cisTEM
173,723 particles
3D classification (D2) - RELION
33007 particles
13.9Å
112919 particles
7.4 Å
26058 particles
11.1Å
3D refinement_ (D2) - RELION
No mask
(4.1Ä)
Mask (3.9Å)
CTF refinement

**Figure S2A**

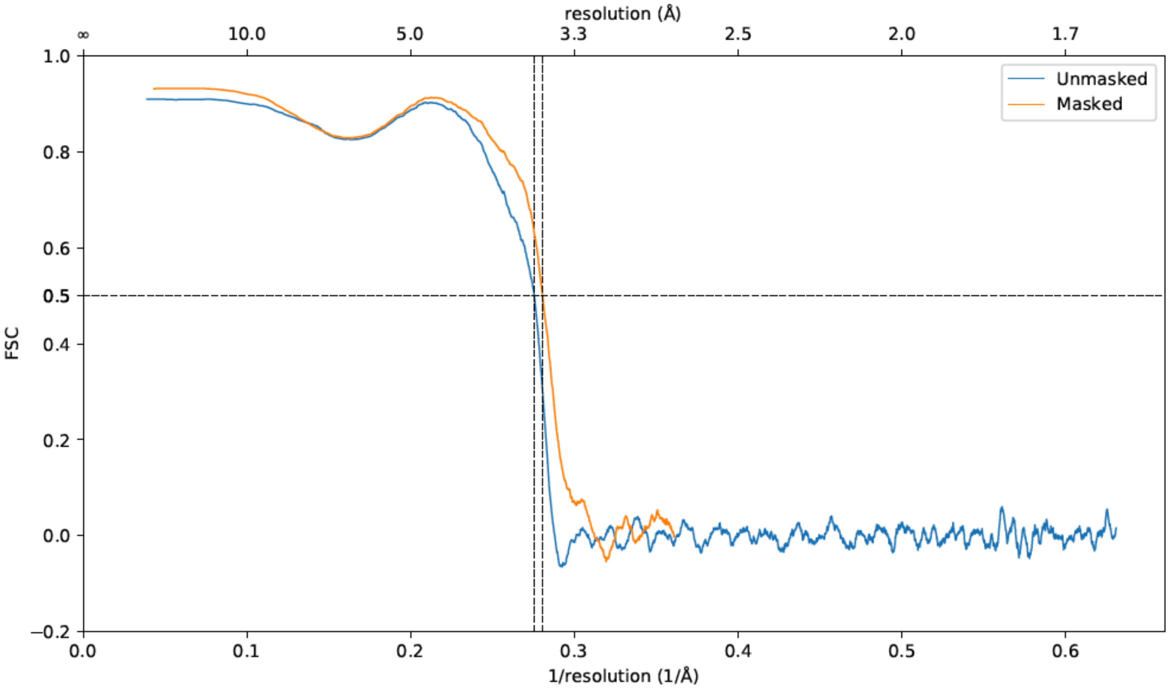


**Figure S2B**

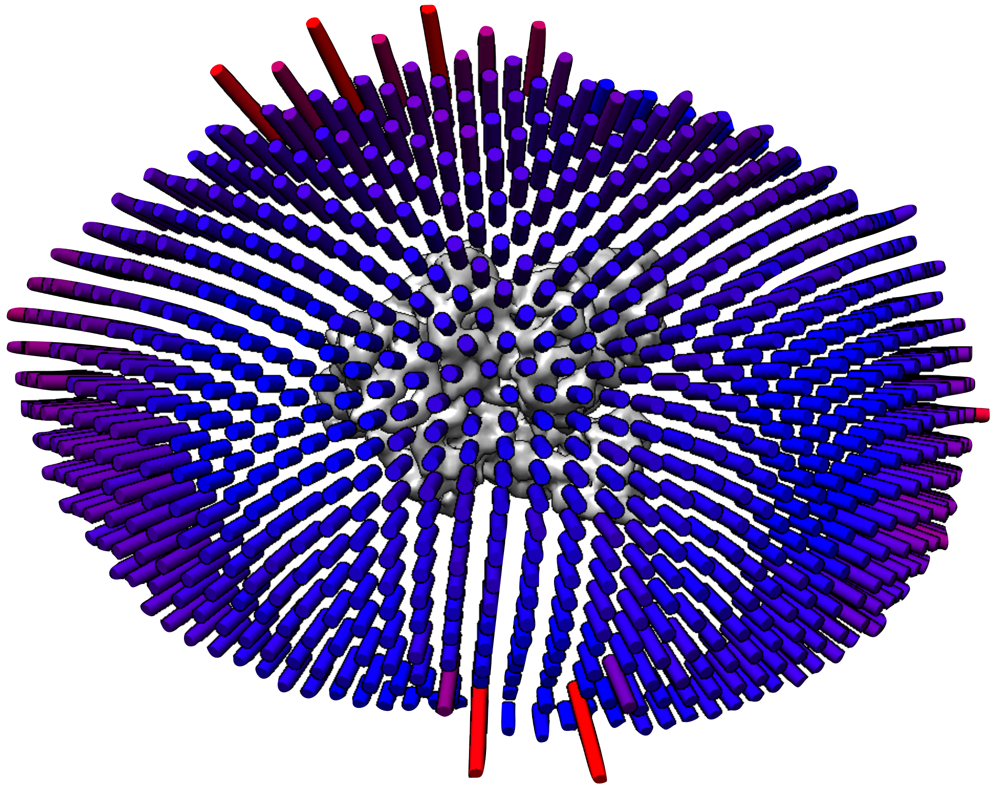

**Figure S2- Map Resolution**: (A) FSC between atomic model and experimental map. (B) Euler angle distribution of particles in final 3D reconstruction.

Figure S3A

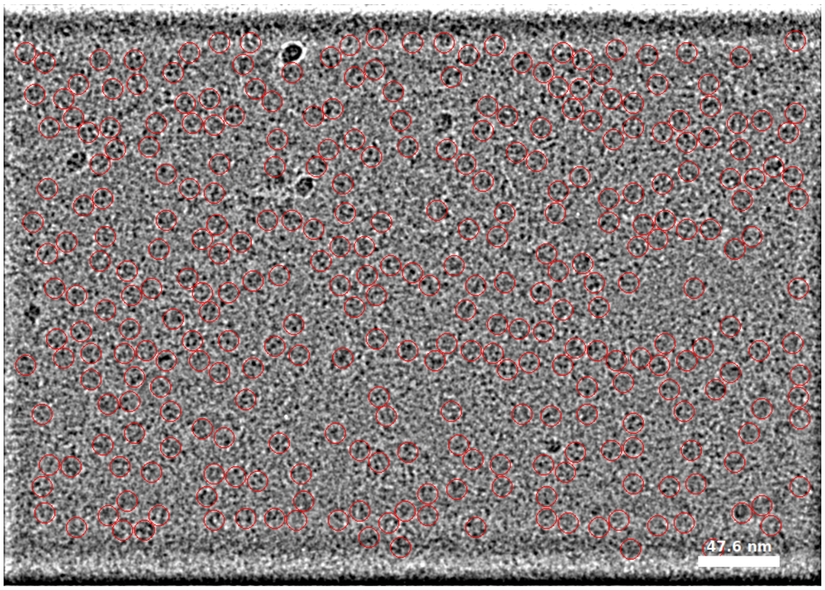


Figure S3B

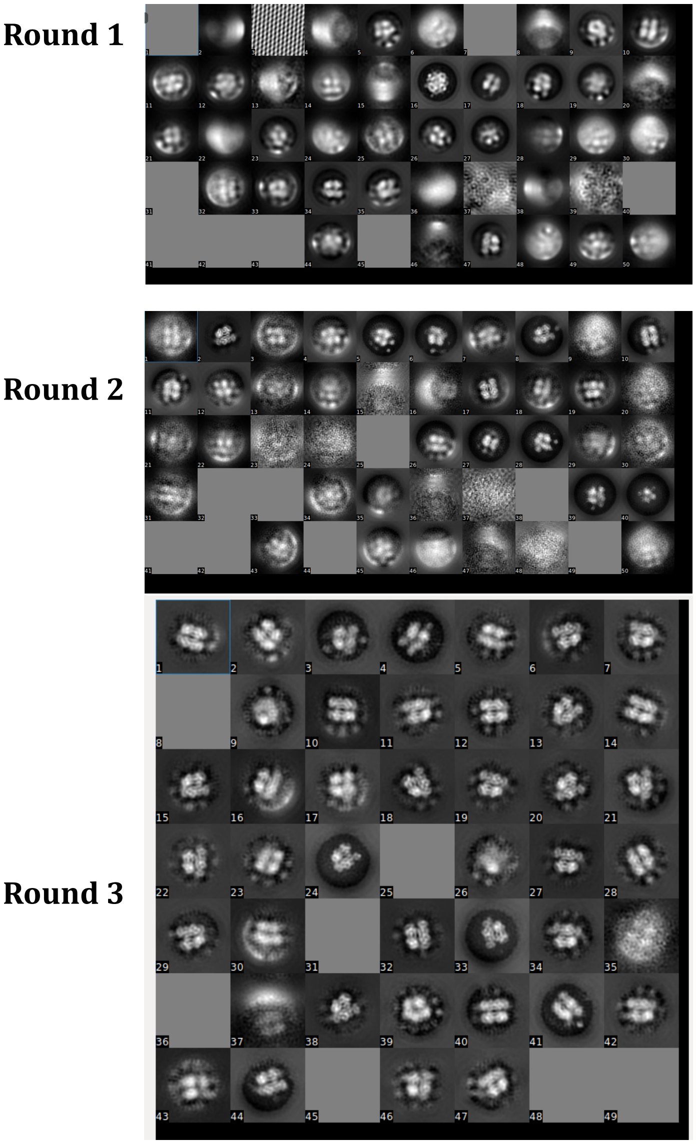

**Figure S3C**

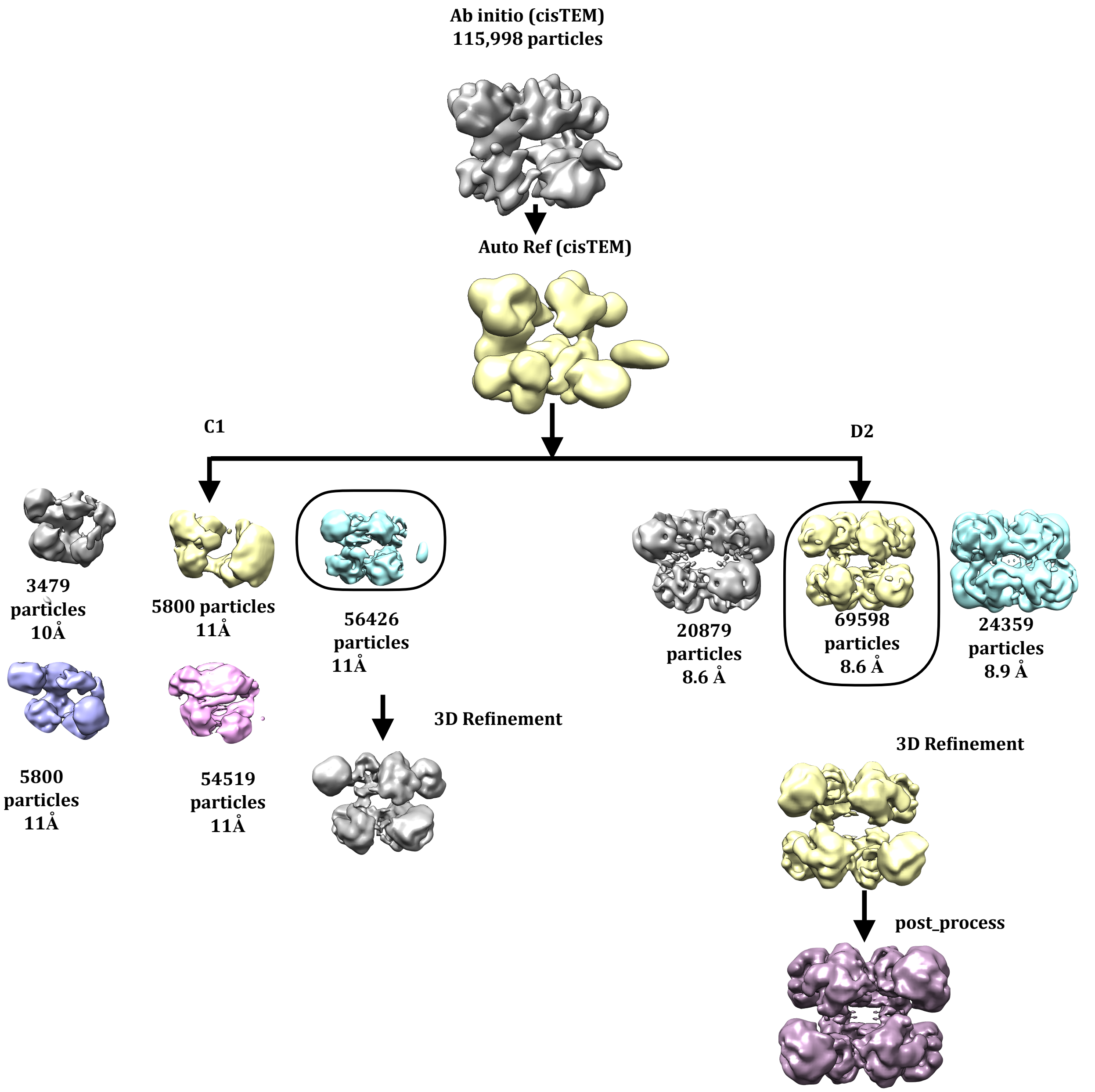


**Figure S3D**

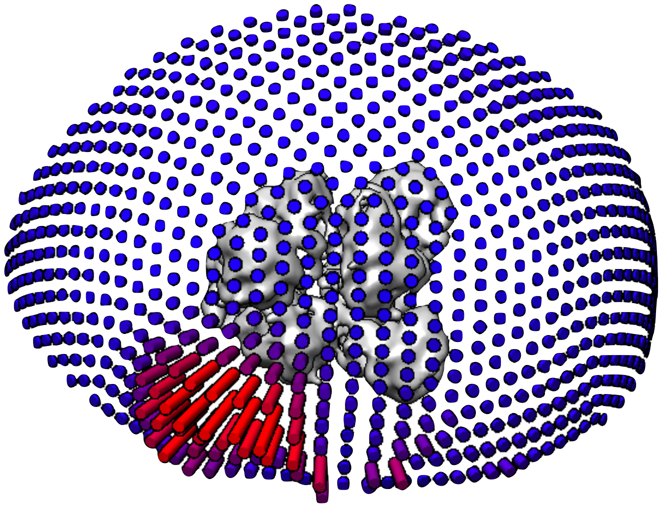

**Figure S3_ CryoEM pipeline of CtBP2$_{31-445}$:** (A) Raw micrograph of CtBP2$_{31-364}$. Particle picking was done in cisTEM. Red circles indicate picked particles. (B) Three round of 2D classification. (C) 3D reconstruction and refinement. (D) Euler angle distribution.

**S4**

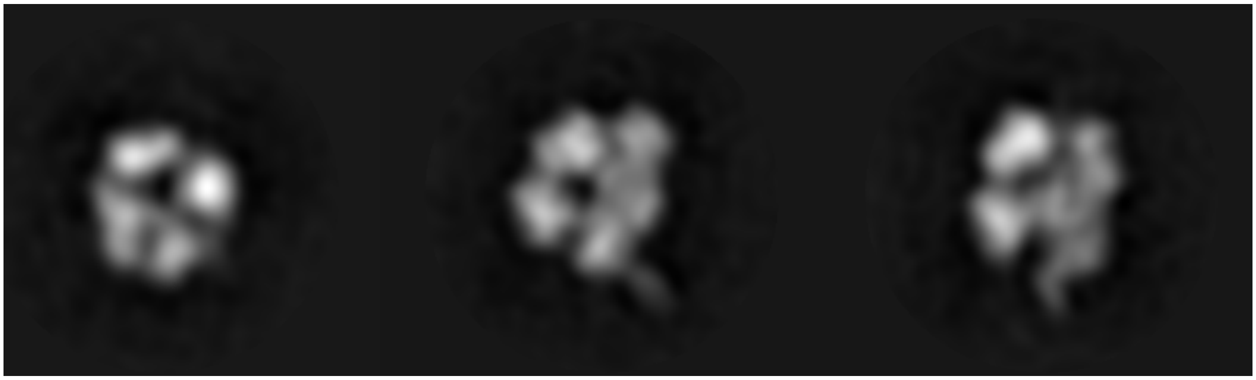

**Figure S4**- Orthogonal slices of the ab initio reconstruction of CtBP2$_{31-364}$ showing

**Figure S5**

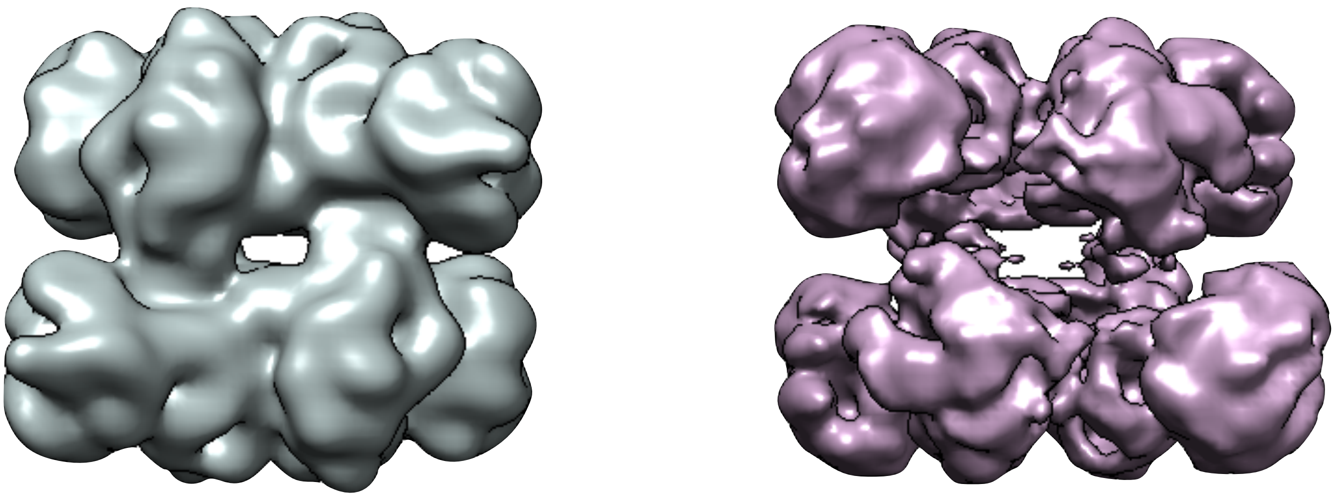

**Figure S5**- Side by side comparison of a low-pass filtered map of CtBP2$_{31\text{-}364}$ and CtBP2$_{31\text{-}445}$.

**Figure S6**

A

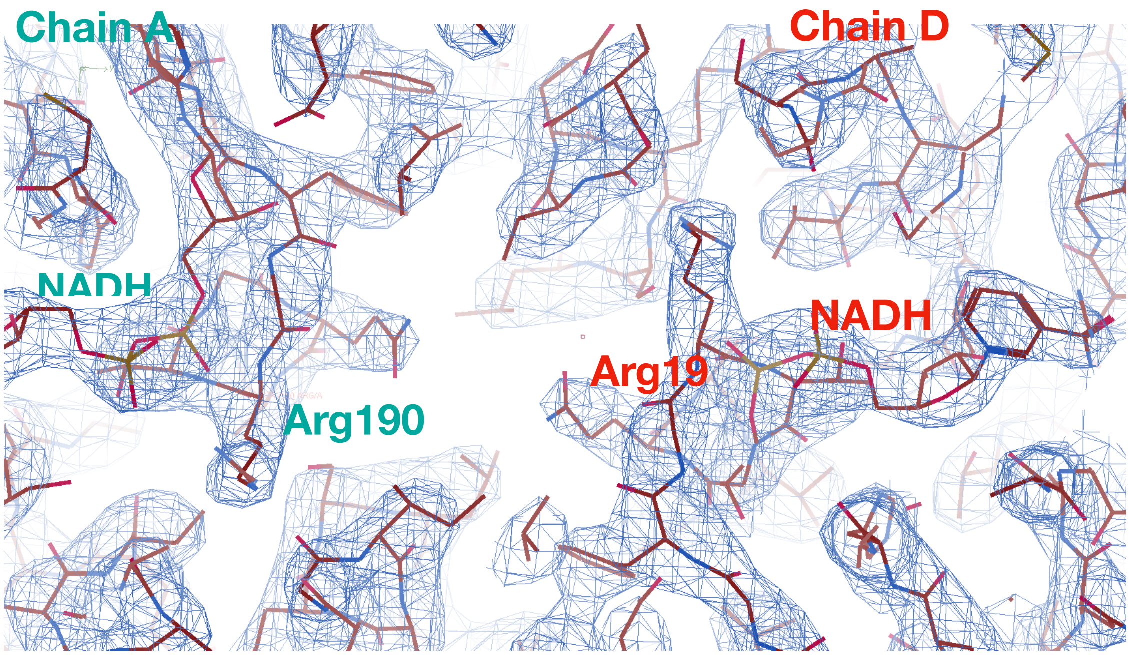


B

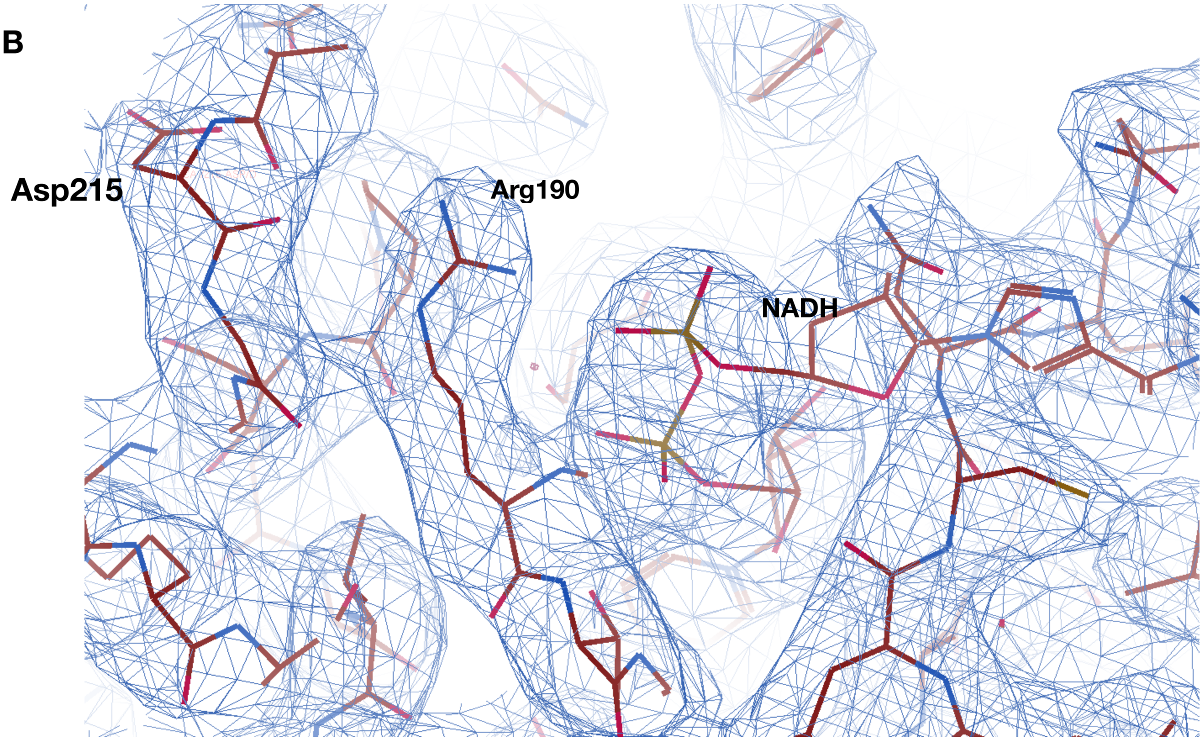


**Figure S6**- Representative density of dimer interface and interactions.

#### Figure S7

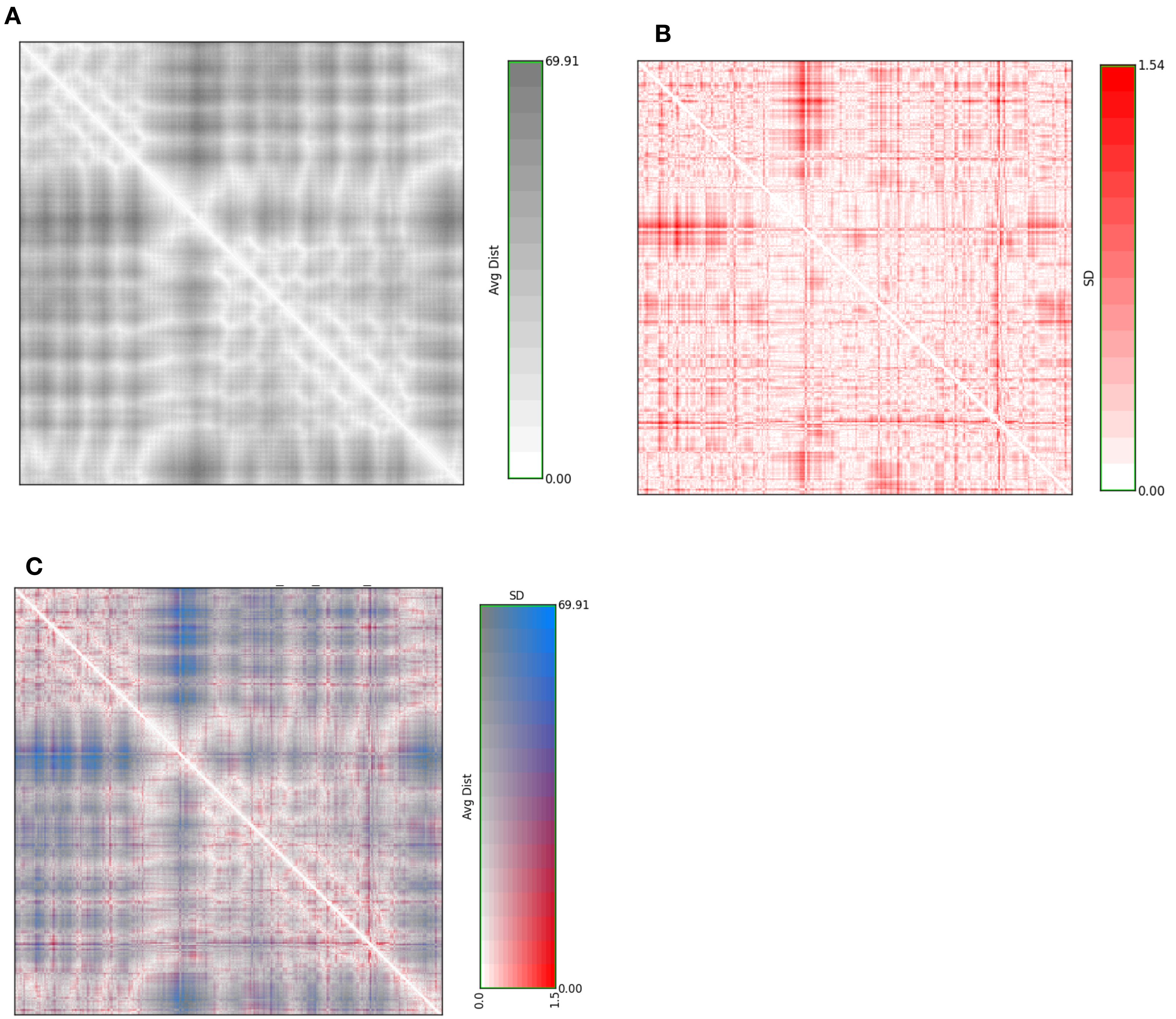


**Figure S7- Cα-Cα distance map analysis of cryoEM and Crystallography structures.** (A) Average distance in Å . (B) Standard deviation in Å. (C) Average distance (Y-axis) and Standard deviation (X-axis) plotted on the same 2D matrix. Distances and deviations calculations were done with residues from all chains.

**Figure S8**

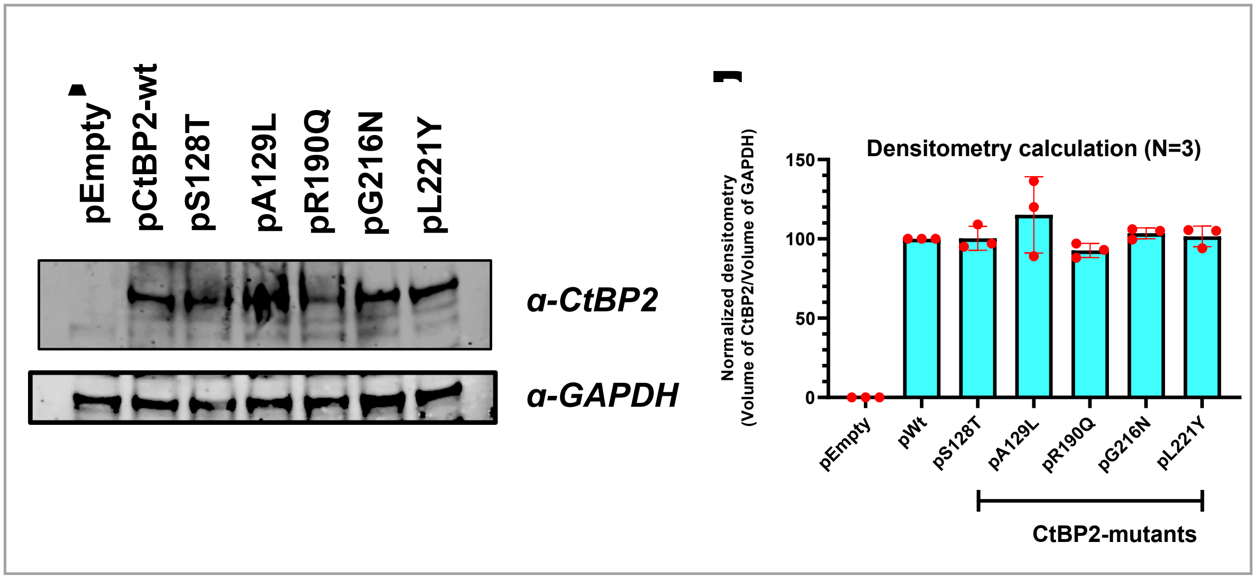


**Figure S8**- **Expression level of CtBP2 variants.** (A) Representative CtBP2 immunoblot of lysates from transfected HCT116; CtBP2(-/-) cells. (B) Mean CtBP2 and CtBP2 variant abundance derived by densiotmetric quantification of CtBP2 immunoblots performed on transfected cell lysates from each replicate. N=3. Red circles indicate individual values from each replicate, and error bars indicate +/- 1 standard deviation.
